# Accounting for red blood cell accessibility reveals distinct invasion strategies in *Plasmodium falciparum* strains

**DOI:** 10.1101/626044

**Authors:** Francisco Cai, Tiffany M. DeSimone, Elsa Hansen, Cameron V. Jennings, Amy K. Bei, Ambroise D. Ahouidi, Souleymane Mboup, Manoj T. Duraisingh, Caroline O. Buckee

## Abstract

The growth of the malaria parasite *Plasmodium falciparum* in human blood causes all clinical manifestations of malaria, a process that begins with the invasion of red blood cells. Parasites enter red blood cells using distinct pairs of parasite ligands and host receptors that define particular invasion pathways. Parasite strains have the capacity to switch between invasion pathways. This flexibility is thought to facilitate immune evasion against particular parasite ligands, but may also reflect the fact that red blood cell surfaces are dynamic and composed of heterogeneous invasion targets. Different host genetic backgrounds affecting red blood cell structure have long been recognized to impact parasite growth *in vivo*, but even within a host, red blood cells undergo dramatic changes in morphology and receptor density as they age. The consequences of these heterogeneities for parasite growth *in vivo* remain unclear. Here, we measured the ability of laboratory strains of *P. falciparum* relying on distinct invasion pathways to enter red blood cells of different ages. We estimated invasion efficiency while accounting for the fact that even if the red blood cells display the appropriate receptors, not all are physically accessible to invading parasites. This approach revealed a tradeoff made by parasites between the fraction of susceptible cells and their invasion rate into them. We were able to distinguish between “specialist” strains exhibiting high invasion rate in fewer cells versus “generalist” strains invading less efficiently into a larger fraction of cells. We developed a mathematical model to predict that infection with a generalist strain would lead to higher peak parasitemias *in vivo* when compared with a specialist strain with similar overall proliferation rate. Thus, the heterogeneous ecology of red blood cells may play a key role in determining the rate of parasite proliferation between different strains of *P. falciparum*.

## Introduction

Proliferation of *Plasmodium falciparum* in human red blood cells underlies all clinical manifestations of malaria, and higher parasite densities in the blood are associated with more severe disease (1, 2). Parasite growth relies on the invasion of red blood cells, mediated by interactions between specific pairs of parasite ligands and host receptors, with each ligand-receptor pair defining a molecular invasion pathway (3). Parasites can switch between invasion pathways via genetic and epigenetic mechanisms; these systems are thought to have evolved in the parasite to evade immune responses against particular ligands (4-7). Redundant invasion pathways may also represent an adaptation to accommodate within-and between-host diversity in red blood cell structure and receptor composition (8). While the importance of red blood cell genetic polymorphisms within human populations in determining parasite growth and disease severity has long been recognized (9-16), the substantial diversity of red blood cells within the host has received less attention. The factors that determine invasion pathway preferences *in vivo*, and their consequences for parasite growth and virulence, remain unclear.

Red blood cells are highly heterogeneous targets for invasion, exhibiting dramatic changes in both the composition and density of potential receptors over the course of their 4-month lifespan. For example, oxidative damage accumulated over this period results in the loss of 10-15% of the cell’s sialic acid (SA) content, reducing the number of glycophorin receptors that can be used by the parasite to enter cells (17-23). Restrictions in receptor availability across differently aged red blood cells are likely to significantly impact the parasite’s *in vivo* proliferation. Indeed, differences in peak parasitemia and virulence between *P. falciparum* and *P. vivax* have been attributed to the restriction of *P. vivax* to invading reticulocytes, the youngest red blood cells (24, 25); by contrast, *P. falciparum* can invade a broader age range of cells, although it too exhibits a preference for younger cells (26). The bloodstream represents a complex and changing ecology, particularly during chronic blood-stage infections in which the distribution of red blood cells may shift over time, as well as in regions where super-infection is common, forcing parasites to compete for resources. As a result, flexible strategies for invasion may be key to enhancing parasite fitness within individual hosts.

The *P. falciparum* genome encodes at least 10 invasion ligands belonging to erythrocyte-binding like (EBL) and reticulocyte-binding like (RBL) protein families (19, 27), each of which binds to a specific red blood cell receptor. Although not all host receptors have been identified, in general, EBL ligands rely on SA-based glycophorin receptors, whereas RBL ligands bind less well-defined receptors that include the recently identified Basigin and Complement Receptor 1 (20, 28). *In vitro,* different *P. falciparum* laboratory strains exhibit distinct preferences for particular invasion pathways (29-31). This flexibility has also been observed in field isolates, with parasite lines from Kenya, Senegal, and the Gambia demonstrating diverse invasion pathway utilization (32-34). Some parasites can switch at high frequency between alternative pathways (Dolan et al., 1990, J Clin Invest). The different members of the RBL and EBL ligands are not essential to invasion in all *P. falciparum* strains, with the PfRh5 parasite ligand being the exception (20, 35), suggesting a functional redundancy in invasion strategies that provides flexibility in the face of immune responses and the heterogeneous ecology of red blood cell receptors.

The importance of different invasion pathways for infection dynamics and parasite growth *in vivo* is poorly understood. Metrics designed to measure the potential for parasite growth in the host using parasites cultured *in vitro* include the commonly measured parasite multiplication rate (PMR), which is analogous to the basic reproduction number (R_0_) in disease ecology, and the selectivity index (SI), which uses the ratio of observed-to-expected frequency of multiply invaded red blood cells to quantify the restriction of invasion to particular subsets of cells. However, both have exhibited inconsistent relationships with peak parasite densities and virulence *in vivo*; PMR calculated from the trajectory of controlled infections in patients with neurosyphilis was shown in the early 20^th^ century to be correlated with peak parasite densities (36). Some studies of natural infections in Southeast Asia have also shown a correlation between PMR and disease severity, and an inverse correlation between SI and severity (24, 37), but African studies have uncovered no such relationships (32, 38).

We propose that variation in parasite proliferation *in vivo* among *P. falciparum* genotypes may reflect diverse ecological strategies – mediated by the use of different invasion pathways – in heterogeneous red blood cell populations. In other words, *P. falciparum* strains of distinct genotype may be specialized with respect to the ability to invade red blood cells of different age, just as different species of *Plasmodium* are. To investigate this, we measured the ability of laboratory-adapted strains of *P. faliciparum* with known differences in invasion pathway utilization to enter red blood cells of different age. We developed a method to jointly estimate the fraction of red blood cells susceptible to invasion and the invasion rate into susceptible red blood cells, to account for the fact that many cells may not be susceptible to invasion simply because they are physically inaccessible to parasites. Our approach revealed different ecological strategies between parasite strains with respect to the age of the red blood cell that are not captured using standard methods. Using a dynamic model of parasite invasion and growth *in vivo*, we predict that these differences would lead to different infection dynamics and peak parasite densities. The complex ecology and dynamics of human blood is therefore an important but under-studied driver of malaria infection dynamics, providing the backdrop for variation in malaria parasite virulence.

## Results

### Accounting for red blood cell accessibility reveals variable age preferences between *P. falciparum* strains

To disentangle the implications of invasion pathway variation and the changing availability of host receptors as red blood cells age, we separated red blood cells into different age fractions (see Methods). We conducted invasion assays into these red blood cell subpopulations with four *P. falciparum* parasite laboratory strains (3D7, Dd2, HB3, and FCR3-SV) that use alternative invasion pathways, characterized by their dependence on SA-containing or chymotrypsin-or trypsin-sensitive receptors (19, 29). In addition, we analyzed the invasion of two strains, Dd2Nm and C2, which are isogenic to Dd2 and 3D7 respectively, with changes in expression of single invasion ligands (increased Rh4 expression in Dd2Nm (39) and loss of Rh2b expression in C2 (40), causing them to shift in their relative dependence on SA for invasion. Invasion assays were plated at the same pre-invasion parasitemia, and the post-invasion parasitemia was measured. To quantify parasite growth while controlling for variability between trials, we calculated the post-invasion parasitemia relative to that of the pooled positive control – we will refer to this measure as the “relative parasitemia”. To estimate parasite selectivity for red blood cell targets, we counted the number of parasites in invaded cells to generate a distribution of multiply infected cells, consistent with previous studies (24). Importantly, we used a standard static assay to improve our ability to quantify multiply infected cells, a key parameter in the measurement of invasion pathway.

The physical inaccessibility of red blood cells in static cultures relative to continuously agitated cultures can significantly alter measures of parasite growth and selectivity (24). Using the Boolean-Poisson model from percolation theory, we estimated that over 97% of red blood cells are likely to be inaccessible to invasion in static culture (see Supplementary Materials for estimates of accessibility). To account for the fact that not all cells are both permissive and physically accessible to invasion, which we will refer to as susceptible, we modeled the distribution of invasion multiplicity using a zero-inflated Poisson model, which allowed us to jointly estimate how many cells pre-invasion were actually susceptible and the invasion rate into those cells (Fig. 1A, Methods). The zero-inflated Poisson model is parameterized by *f* and *λ*, where *f* is the probability that the data are generated from a Poisson process with rate *λ*, and 1 − *f* is the probability of a structural zero. In our biological context, *f* is the fraction of red blood cells available for invasion (fraction susceptible), and *λ* is the invasion rate. Note that the original SI (24) assumed the Poisson as a null model, with all cells being susceptible to invasion. We will refer to invasion rate estimates based on the zero-inflated Poisson model as “adjusted” estimates and those based on a Poisson model as “unadjusted”. We compared the two models using a likelihood ratio test, and for 169 (89%) of the 190 invasion assays performed, the zero-inflated Poisson model gave a significantly better fit than the Poisson model, assuming an overall significance level of 0.05 with Bonferroni correction (Fig. S1).

**Fig. 1.**
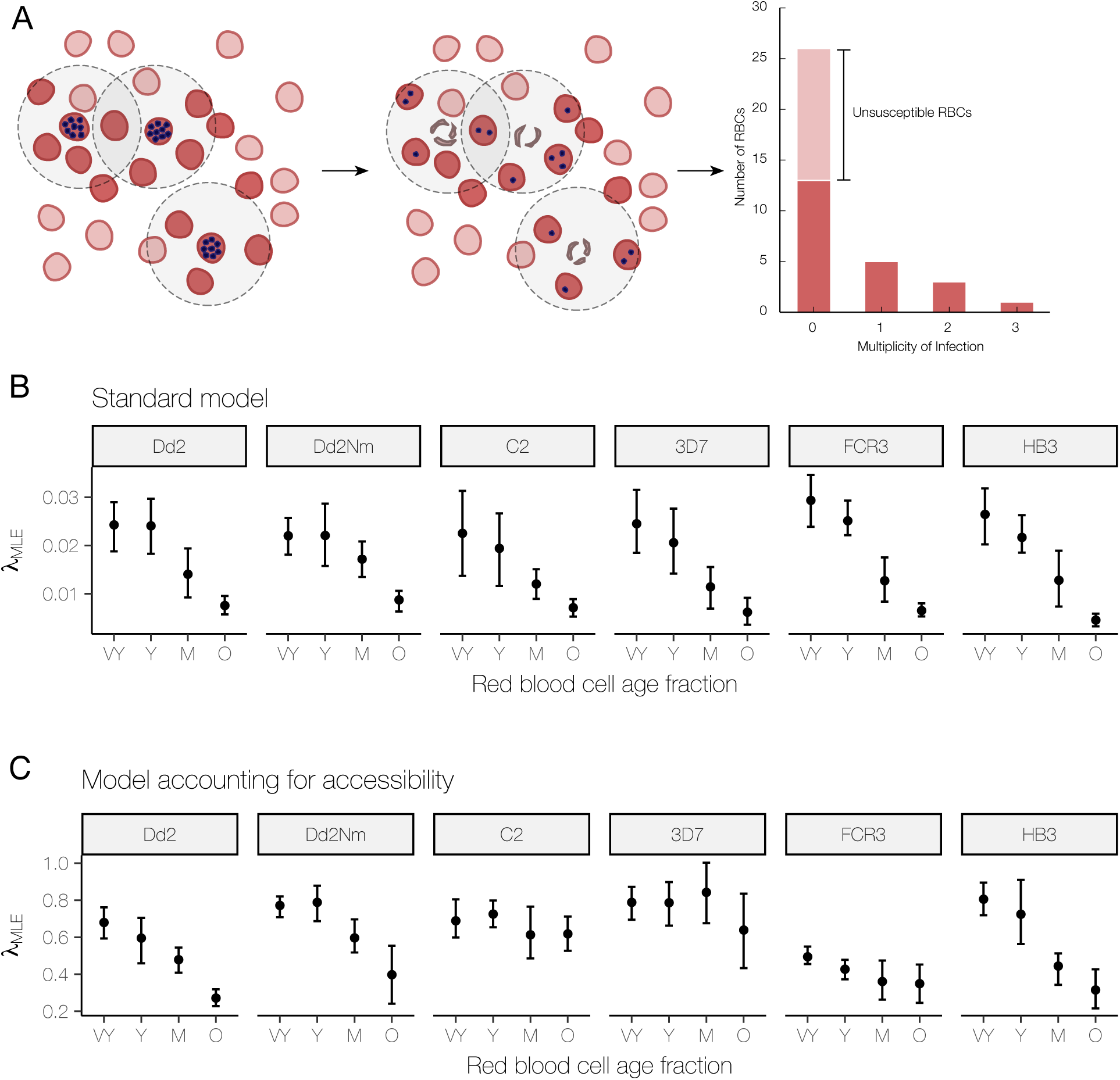
Accounting for accessibility reveals different responses to red blood cell age. **(A)** Pre-invasion *(left)*: Red blood cells susceptible to invasion (darker red) are both accessible – within the burst radius of a schizont (blue dots represent merozoites) – and permissive to invasion. Unsusceptible red blood cells (light red) may be too far from a schizont or within the burst radius but not permissive. Post-invasion *(middle)*: After schizonts burst, susceptible red blood cells may escape infection or be newly infected with 1 or more merozoites. Distribution of infection multiplicity *(right)*: Post-invasion, we count the number of red blood cells infected with 0, 1, 2, and 3 merozoites to generate a distribution of infection multiplicity. A Poisson rate can be estimated using the distribution that includes all red blood cells or a distribution that considers only susceptible red blood cells (darker red bars). **(B)** Maximum-likelihood estimates of the unadjusted invasion rate (***λ***_***MLE***_) using the standard model. The distributions (mean with 95% bootstrapped CI) of invasion rate estimates (N = 5 to 8 trials), organized by the parasite strain (panel title) and red blood cell age fraction (x-axis, VY = very young, Y = young, M = medium, O = old) used in the invasion assay. **(C)** Maximum-likelihood estimates of the adjusted invasion rate (***λ***_***MLE***_) using the model accounting for accessibility. The distributions are presented in the same way as in (B).

Once we accounted for restricted physical access to host targets, we measured increased invasion rates and substantial differences between strains in their ability to invade red blood cells of different ages. The unadjusted estimates of the invasion rate declined with increasing red blood cell age for all laboratory strains (Fig. 1B), whereas the adjusted estimates showed that two laboratory strains, C2 and 3D7, maintained a constant invasion rate with increasing red blood cell age (Fig. 1C). This suggests that the decreasing trends for these strains using the standard approach were caused by the reduced number of cells being available for invasion in the older fractions rather than reduced efficiency of invasion into them. The fraction susceptible, *f*, has a natural interpretation as a measure of selectivity, with a lower fraction susceptible indicative of a more selective invasion process. Our fraction susceptible estimates are related to the SI – which is essentially a measure of how poorly the Poisson model fits the observed data – and we found a strong negative correlation between the fraction susceptible and the SI (Spearman’s rank correlation = −0.95, N = 190). Unlike the SI, however, the fraction susceptible is biologically interpretable and amenable to maximum likelihood statistics (providing confidence intervals) and is thus a rigorous alternative to quantifying selectivity in parasite growth.

### *P. falciparum* strains with similar multiplication rates exhibit distinct invasion strategies with respect to red blood cell age

We mapped *λ* (invasion efficiency) and *f* (fraction susceptible) for four lab strains – Dd2, 3D7, FCR3, and HB3 – with known differences in invasion phenotype, but similar PMR in whole blood. Although the relative parasitemia for a given red blood cell fraction was similar between strains (Fig. S2), this was achieved with different tradeoffs between fraction susceptible and invasion rate (Fig. 2A). We propose that these differences may reflect different ecological strategies for invading heterogeneous red blood cells, which we place along the spectrum from generalist to specialist (Fig. 2C). For example, FCR3 has a generalist invasion strategy: a higher fraction of red blood cells is susceptible to invasion, but the invasion rate into these cells is lower. On the other hand, 3D7 appears to have a more specialist strategy: a lower fraction of red blood cells is susceptible, but the invasion rate into the susceptible fraction is higher. Dd2 and HB3 exhibit an intermediate strategy. We also compared two pairs of isogenic laboratory strains, Dd2/Dd2Nm and C2/3D7, where the strains in each pair differ by a single point mutation that renders them SA-dependent/independent, respectively. Fig. 2B illustrates that within each pair, the SA-independent strain displayed a more specialist phenotype (i.e. for each red blood cell age fraction, the SA-independent strain had a lower fraction susceptible, but a higher invasion rate than the corresponding SA-dependent strain).

**Fig. 2.**
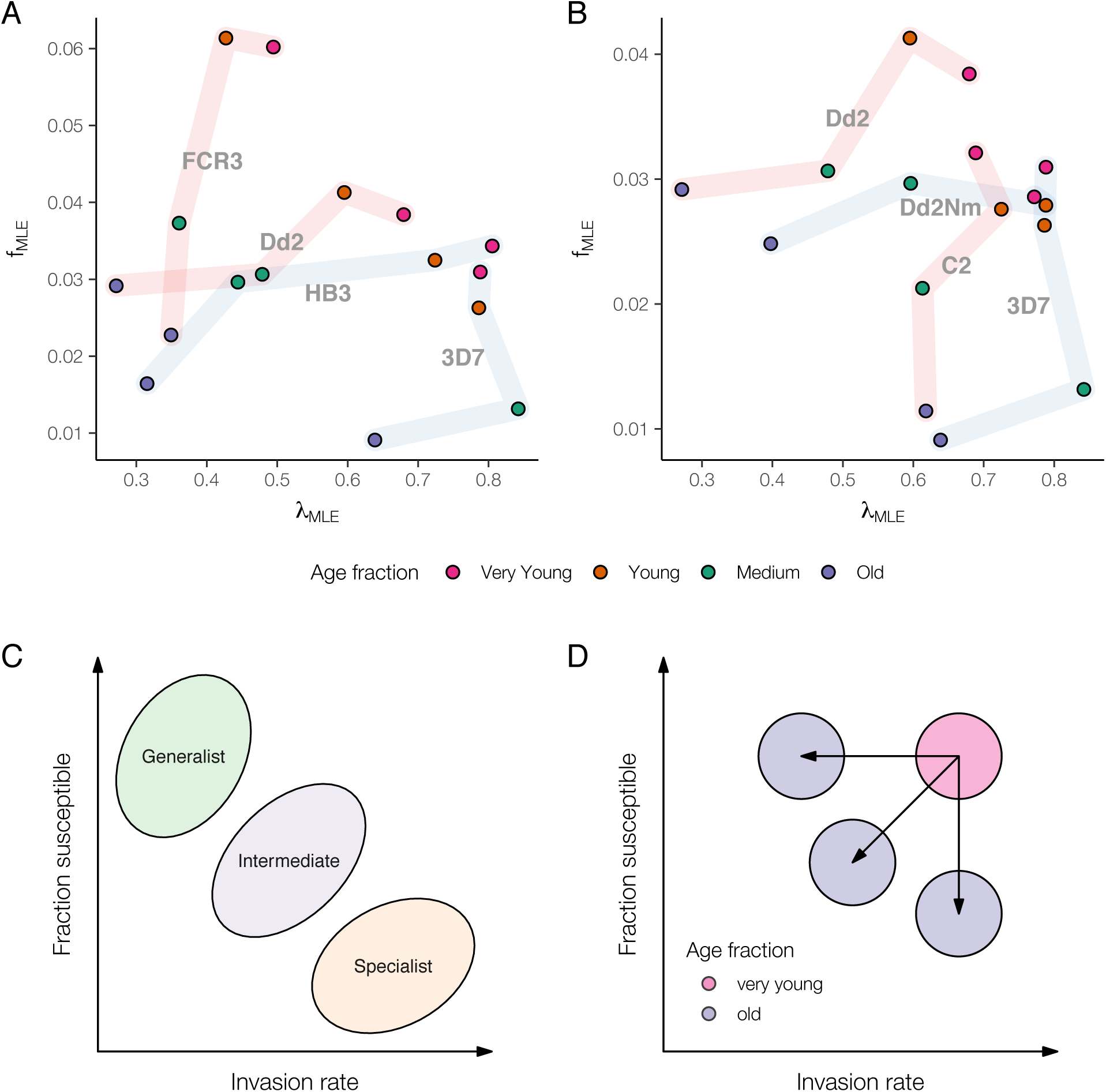
**(A, B)** Modeling susceptibility reveals distinct invasion strategies. Each point denotes the maximum likelihood estimate of the fraction susceptible ***f***_***MLE***_ (y-axis) and ***λ***_***MLE***_ (x-axis), colored by the age fraction and labelled by the parasite strain used in the invasion assay. Points for the same strain are connected by a light band in order of increasing red blood cell age. A light red band indicates that the strain is sialic-acid dependent (FCR3, Dd2, C2); blue, sialic-acid independent (HB3, 3D7, Dd2Nm). **(B)** shows four laboratory strains: FCR3, Dd2, HB3, and 3D7. **(C)** shows two isogenic strain pairs, Dd2/Dd2Nm and C2/3D7, where the strains in each pair differ by a single point mutation that is associated with their dependence on sialic-acid receptors. **(C, D)** Conceptualizing invasion strategies and response to red blood cell aging. **(C)** Parasites with similar PMR can be distinguished in terms of how they tradeoff between invasion rate and fraction susceptible, with generalists—low invasion rate into a high fraction susceptible—and specialists—high invasion rate into a low fraction susceptible—on either end of the spectrum. **(D)** Strains can be further distinguished by their response to aging red blood cells— whether they experience a decline in invasion rate (horizontal arrow), a decline in fraction susceptible (vertical arrow), or a combination of both (diagonal arrow).

A further distinction can be made with respect to red blood cell aging. Although relative parasitemia declines in older red blood cells for every strain (Fig. S1), the decline occurs in different ways: FCR3 and 3D7 experience a steep drop in fraction susceptible with relatively little change in invasion rate, whereas Dd2 and HB3 experience a smaller decrease in fraction susceptible, but a larger decrease in invasion rate (Fig. 2A). The paired isogenic strains had similar responses to red blood cell aging: Dd2 and Dd2Nm had a modest decrease in fraction susceptible, but a larger decrease in invasion rate, whereas C2 and 3D7 had a substantial decrease in fraction susceptible, but a relatively small decrease in invasion rate (Fig. 2B). Figure 2D illustrates a conceptual model of these differential responses to cell aging. Of note, *ex vivo* assays of field isolates from clinical malaria cases in Senegal showed variation across these gradients, both in terms of overall strategy (Fig. S3A) and in their response to red blood cell aging (Fig. S3B).

### Diverse invasion strategies generate important differences in within-host dynamics

To predict the effects of different invasion strategies on within-host dynamics, we used a simple compartmental model with three red blood cell compartments (susceptible, non-susceptible, and infected) and one merozoite compartment (see Methods). We compared hypothetical strains that achieve the same PMR through different combinations of fraction susceptible (denoted *f*) and invasion efficiency (denoted *p* and defined as the probability that a parasite successfully invaded after coming into contact with a permissive red blood cell). For a fixed PMR, the fraction susceptible and invasion efficiency are inversely proportional (Fig. 3A). We included a specialist strain (*f* = 0.1, *p* = 0.83), a generalist strain (*f* = 0.5, *p* = 0.17), and an intermediate strain (*f* = 0.25, *p* = 0.33). Despite all three strains having the same PMR, the generalist strain had the highest peak parasitemia, and the specialist strain had the lowest (Fig. 3B). Thus, we predict that PMR may not provide information about the proliferation potential of a parasite strain and that – consistent with some, but not all previous observations of SI – specialization for particular red blood cell age subpopulations may lead to lower growth rates *in vivo*.

**Fig. 3.**
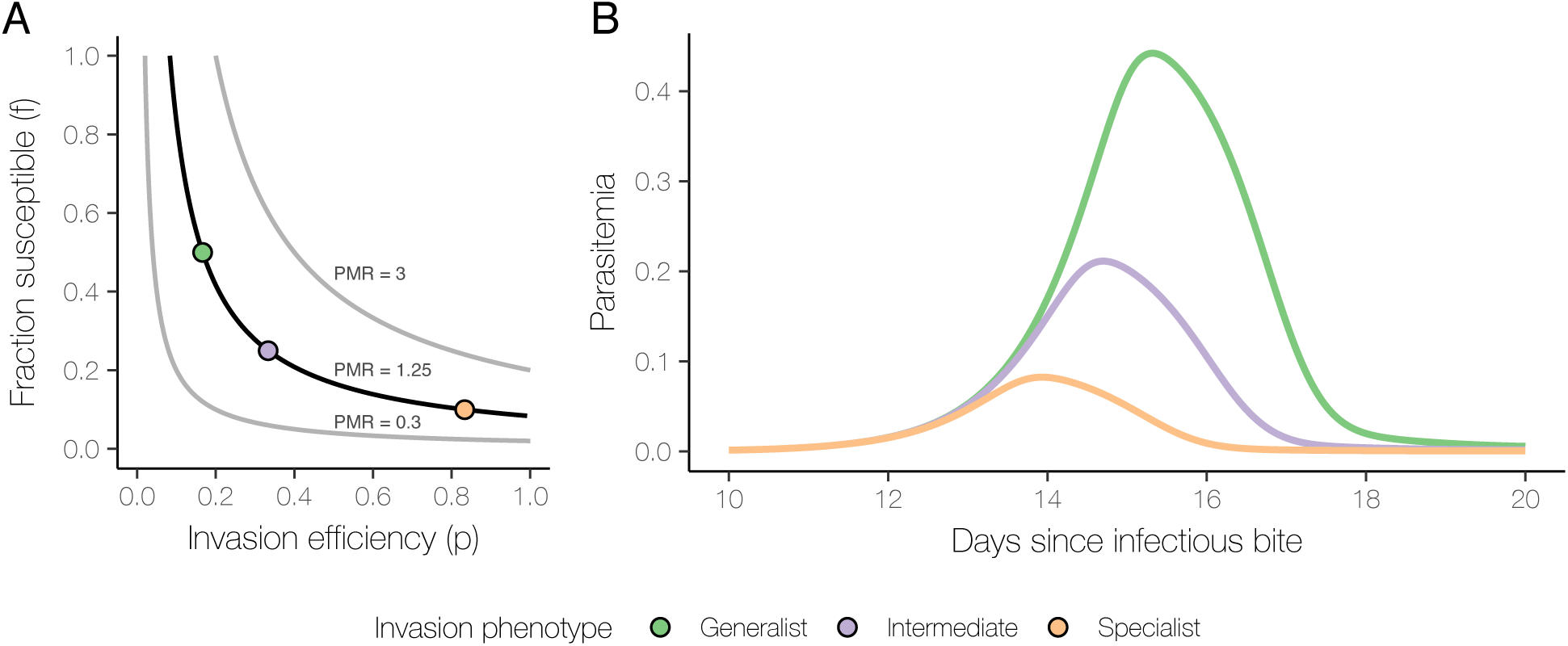
Simulated in vivo invasion dynamics of different invasion strategies. **(A)** The three curves show the inversely proportional relationship between fraction susceptible and invasion efficiency for different fixed values of PMR: 0.3, 1.25, or 3. For our simulations, we considered three invasion phenotypes corresponding to a PMR of 1.25 (black curve): a generalist phenotype (orange point: ***f*** = 0.5, ***p*** = 0.17); an intermediate phenotype (purple point: ***f*** = 0.25, ***p*** = 0.33); and a generalist phenotype (green point: ***f*** = 0.1, ***p*** = 0.83). **(B)** Simulated parasitemia during infection with the specialist (orange), intermediate (purple), or generalist (green) strain for the period 10-20 days after the infectious bite.

## Discussion

Using a new approach to understand the nuances of parasite invasion *in vitro*, we have shown that laboratory lines that would have otherwise been indistinguishable using standard approaches exhibit distinct invasion strategies with respect to the same red blood cell age fraction. Based on our findings, we propose that in addition to pressures from the immune system (41,42), variations in invasion pathway utilization across *P. falciparum* strains may represent an adaptation to the heterogeneous and dynamic ecological landscape of the host blood stream.

Susceptibility to invasion requires that red blood cells be physically accessible to merozoites, which are non-motile, as well as permissive (i.e. expressing receptors required by the parasite). Fitting a Poisson distribution assumes that all red blood cells are equally susceptible and that invasion occurs randomly. In reality, however, a number of factors, such as immunity, rosetting, and differential permissiveness of red blood cells, cause deviations from this model. The SI has been used by some researchers to measure heterogeneous invasion *in vitro* (24, 37, 38), but is insensitive to the distinction between physical inaccessibility and non-permissiveness. To address this issue, we used a zero-inflated Poisson model. This is particularly useful to *in vitro* static cultures, where experimental conditions determine the fraction of cells that are physically accessible to the parasite. Under a zero-inflated Poisson model, the estimated fraction susceptible is indicative of the relative permissiveness of different subpopulations of red blood cells to different parasite strains.

By disentangling these different aspects of invasion, we were able to distinguish between generalist and specialist *P. falciparum* strains. The former is able to invade a larger fraction of red blood cells, but at a lower rate, whereas the latter is able to invade a smaller fraction of red blood cells, but at a higher rate. We have found that even with a pooling of all red blood cell age fractions, the difference between a generalist and a specialist strain remains. Using a compartmental model, we have shown that this has consequences *in vivo*: in comparison with a specialist strain with the same PMR, a generalist strain is likely to exhibit higher parasite densities, which may in turn affect disease severity. This is supported by the work of Lim et al., which suggested that adaption of *P. knowlesi* to a broader age range of red blood cells is the likely the cause of higher parasitemia *in vivo*; this adaptation was found to persist for many generations in parasites cultured *in vitro* (43). Conversely, modeling work by Kerlin et al. has shown that a restricted age range of susceptible red blood cells is a credible mechanism by which parasites may self-regulate proliferation (44). McQueen et al. have also reported on the importance of red blood cell age in predicting infection dynamics within a host (45). Of note, the impact of the heterogeneous susceptibility of cells for infection dynamics is paralleled in models of an infectious disease spreading through an age-structured population of hosts, where epidemic trajectory depends not on the R_0_ overall, but on the R_0_ in different subgroups of the population (46).

Our findings not only have implications for how we measure and understand the relationship between invasion phenotypes and virulence, but also how we predict the impact of the skewing of red blood cell age structure due to anemia and measure growth parameters *in vitro*. Our adjusted methods for measuring these aspects of parasite invasion therefore provide a new tool to assess the strategies used among field isolates in different clinical and transmission settings.

## Materials and Methods

### Density-dependent fractionation of red blood cells

Red blood cells of different ages were fractionated by exploiting density differences among red blood cell subpopulations using a discontinuous Percoll gradient as previously described (47-49). In brief, blood was incubated at 37°C for 30 hours prior to fractionation (50). Five Percoll solutions of different concentrations (40%, 45%, 50%, 53.5%, and 65%) were diluted in un-supplemented RPMI. Two mL of each dilution were overlaid into a 15 mL Falcon tube from most dense (65%) to least dense (40%). 500 μL of a 30% hematocrit suspension of washed O^+^ blood cells was then added. The gradient was centrifuged for 12 min at 1200 x g, at 22°C. Four prominent bands corresponding to red blood cells of different ages (very young, young, medium and old) were resolved, with the densest (oldest) cells settling into the bottommost layer. Although the relative abundance of each age fraction varied by blood donor, the approximate proportions of red blood cell fractions were as follows: 40%, 30%, 20%, and 10% for very young, young, medium, and old, respectively.

### ADVIA analysis

An ADVIA hematological analyzer (Bayer) was used to validate the efficacy of fractionation of whole blood into discrete age groups. Following Percoll fractionation, each of the red blood cell subpopulations was washed and re-suspended in 1x PBS. Measurements were taken for each age fraction, as well as a positive control consisting of Percoll-fractionated blood that was subsequently re-pooled. Older cells were associated with decreased cell size (mean corpuscular volume [MCV]) and increased density (mean corpuscular hemoglobin concentration [MCHC]), demonstrating the isolation of distinct red blood cell subpopulations (52).

### Age-dependent invasion assay

Age-dependent invasion efficiencies were measured using standard invasion assays. Invasion assays were performed on sorbitol-synchronized, ring-stage parasites plated at a final parasitemia of 0.7% and a hematocrit of 2%. Parasitized donor cells were added to an equal volume of age-fractionated (very young, young, medium, and old) red blood cells (receptor cells) in a 96-well plate. Donor cells were treated with both neuraminidase (Calbiochem, 66 mU/ml) and chymotrypsin (Worthington Biochemicals, 1 mg/ml) for 1 hour at 37°C. This enzyme treatment cleaves the majority of known red blood cell receptors and renders these cells refractory to parasite invasion, ensuring that only age-fractionated receptor cells are susceptible to invasion. The negative control was composed of receptor and donor cells that were enzyme-treated with neuraminidase and chymotrypsin. The positive control consisted of Percoll-fractionated cells that were subsequently re-pooled (and therefore representative of whole blood), but subjected to the same manipulations as each of the individual red blood cell fractions. Samples were plated in triplicate and incubated at 37°C for 48 hours until parasite re-invasion. Blood smears were then made and parasitemia determined microscopically. Assays were repeated a minimum of five times.

### Statistical analysis of invasion data

Parasitemia was calculated for each assay and divided by the values in the positive control to account for variation between trials to yield the “relative parasitemia”. Since the pre-invasion parasitemia was fixed, the post-invasion parasitemia was proportional to the PMR, and the relative parasitemia was equivalent to the relative PMR (using the positive control as the reference). With relative parasitemia as the outcome variable, we examined the effect of red blood cell age on each strain and the effect of parasite strain within each age group using the non-parametric Kruskal-Wallis test. To account for multiple tests across strata of different red blood cell age groups/parasite strains, the Bonferroni correction was applied. Significant Kruskal-Wallis test results were followed by a post-hoc Dunn’s test to examine pairwise differences between groups. A Type I error rate of 0.05 was used to determine statistical significance. The analysis was performed using R 3.3.3 and the R package dunn.test (53, 54). All six strains exhibited heterogeneity across the different age fractions (Kruskal-Wallis rank sum test, p < 0.01 for all strains), with lower relative parasitemia in older age fractions (Fig. S4). However, within each of the four age fractions, no significant heterogeneity was detected between strains (Kruskal-Wallis rank sum test, p = 0.35, very young; p = 0.94, young; p = 0.43, medium; 0.07, old, Fig. 2).

### Zero-inflated Poisson model of infection multiplicity

In each invasion assay, the number of parasites in each red blood cell (i.e. its multiplicity of infection [MOI]) was determined by microscopy for roughly 300 infected cells. For each MOI, we counted the number of red blood cells with that MOI. The maximum MOI observed was 5. The number of uninfected red blood cells (i.e. having an MOI of 0), was estimated from parasitemia. These counts were used to fit a zero-inflated Poisson model parameterized by *f* and *λ*, where *f* is the probability that the data were generated from a Poisson process with rate *λ*, and 1 − *f* is the probability of a structural zero. Maximum likelihood estimates of the parameters were calculated. A Poisson model with no zero inflation (i.e. *f* = 1) was also fit to the data using maximum likelihood estimation, and the likelihood ratio test was used to compare the zero-inflated and non-zero-inflated models.

### Within-host infection dynamics compartmental model

A continuous-time compartmental model was used to simulate within-host infection dynamics. Four compartments were used: three to track the numbers of susceptible, unsusceptible, and infected red blood cells, and one to track the number of merozoites. The rate of production of new red blood cells was chosen to maintain 20-30 trillion red blood cells in circulation absent infection (55), given a red blood cell lifespan of 110 days (56). Of the new red blood cells, a constant fraction entered as susceptible red blood cells. We assumed that at the start of infection, inoculation with sporozoites resulted in 15 liver-stage parasites, each of which released 40,000 merozoites into circulation (57-60). For simplicity, the rate of infection assumed merozoites contacted a susceptible red blood cell every 10 minutes, resulting in successful invasion with a certain probability that varied between simulations. An infected red blood cell has a lifespan of 2 days, corresponding to the duration of schizogony (61, 62). The fraction of red blood cells susceptible and the probability of infection given contact were varied between simulations.

## Supplemental Information

### Within-host Infection Dynamics Compartmental Model Additional Details

A continuous-time compartmental model was used to simulate within-host infection dynamics. The basic model consisted of three red blood cell compartments – susceptible, unsusceptible, and infected (denoted *S, R*, and *I*, respectively) – and one merozoite compartment, denoted *Z*. The corresponding system of ordinary differential equations (ODEs) is:

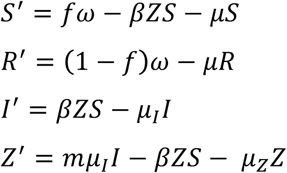

*ω* is the rate of production of new red blood cells, of which fraction *f* are susceptible to invasion. *μ* is the death rate shared by susceptible and unsusceptible red blood cells; *μ*_*I*_ is the death rate for infected red blood cells; *μ*_*Z*_ is for merozoites. *β* is the rate of new infections per susceptible red blood cell per merozoite. *m* is the number of merozoites released from an infected red blood cell when it ruptures. Parameter values are listed in Supplemental Table 1. For the basic model, the waiting time in each compartment followed an exponential distribution. To more realistically model the lifespan of uninfected and infected red blood cells (approximately 110 days and 2 days, respectively), each red blood cell compartment was replaced with a series of successive compartments, so that the total time spent in a compartment series followed a gamma distribution (63). *n* compartments was used to replace the uninfected compartments, and *n*_*I*_ compartments was used to replace the infected compartment, resulting in a modified system of ODEs used in our simulations:

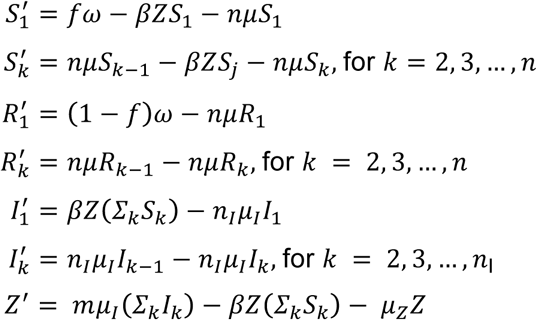

To choose parameter values given a desired PMR, we used the relationship PMR *≅ mfp*, where *m* and *f* are as described above, and *p* is the probability of invasion given contact between a merozoite and a susceptible red blood cell. This is a reasonable approximation of PMR when there are relatively few multiply-infected red blood cells. We then calculated *β* as *p*/*τ*, where *τ* is the length of time needed for the merozoite to complete invasion. Values of *m* and *τ* were determined from the literature. For a fixed PMR, choosing a value for *f* determines the value of *p*, which in turn determines the corresponding value of *β* to be used in the simulation.

For each set of parameter values, the model was simulated for 500 days without merozoites to reach equilibrium, after which merozoites were introduced to the model. We assumed that infection began with 15 infected hepatocytes releasing 40,000 merozoites each, and we continued the simulation for 25 additional days, long enough to observe the first peak parasitemia (55-58, 64, 65).

### Estimation of Physical Accessibility using a Boolean-Poisson Model

We estimated the fraction of red blood cells accessible to parasite invasion using the Boolean-Poisson model from continuum percolation theory. To use this model, we assumed that the locations of red blood cells and schizonts were random (i.e. can be modelled with a spatial Poisson processes) and that when a schizont burst, the released merozoites are able to access red blood cells within a sphere of fixed radius (the burst radius) centered on the location of the bursting schizont. The expected volume fraction accessible by merozoites is then given by

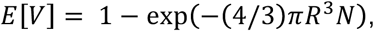

where *R* is the burst radius, and *N* is the average number of schizonts per unit volume, which can be calculated as the product of red blood cell count (number of red blood cells per unit volume) and schizontemia (number of schizonts per red blood cell, denoted *s*). The red blood cell count itself can then be decomposed as the hematocrit (volume fraction occupied by red blood cells, denoted *h*) divided by the MCV. For simplicity, we assumed that red blood cells are spherical with radius *r*. We can rewrite the number of schizonts in terms of the hematocrit, MCV, and schizontemia and assuming that red blood cells are roughly spherical:

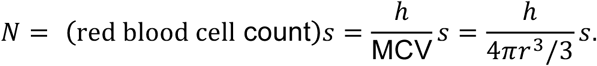

Substituting this expression into our previous equation for the expected volume fraction gives

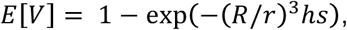

where we have written the expected volume fraction accessible as a function of the burst radius (relative to the red blood cell radius), hematocrit, and schizontemia. Finally, we assumed that red blood cells are distributed randomly in space, so the expected fraction of volume accessible is also the fraction of red blood cells accessible. Crick et al. estimated the dispersion radius of a merozoite group relative to the schizont mean radius to be 1.5 (64). Taking that value to be *R/r* and using the hematocrit and schizontemia from our invasion assays, 2% and 0.7% respectively, we estimated a fraction accessible of less than 0.1%. However, in static culture, there is sedimentation of red blood cells, so the effective hematocrit could be much higher than 2%. Even assuming a hematocrit of 100% (keeping the same values for *R/r* and schizontemia), the fraction accessible increases to only 2.3%.

---

**Supplemental Table 1.** Parameters of the within-host compartmental model.

| Symbol | Description | Value | Reference |
| --- | --- | --- | --- |
| Biological Parameters |  |  |  |
| $\omega$ | Rate of hematopoiesis | 0.25 trillion red blood cells (RBC) / day | Rate chosen to maintain 20-30 trillion circulating RBCs (53) |
| $f$ | Fraction of RBCs susceptible to invasion | 0.1, 0.25, 0.5 | — |
| $p$ | Probability of invasion given contact with susceptible RBC | 0.83, 0.33, 0.17 | For a fixed PMR, value determined by $f$ |
| $\tau$ | Duration needed for invasion, given contact has occurred | 5 min | (65) |
| $\beta$ | Rate of invasion per contact with susceptible RBC | 250, 100, 50 day<sup>-1</sup> | Derived as $p/\tau$ |
| $\mu$ | Death rate for uninfected RBCs | 1/110 day<sup>-1</sup> | (54) |
| $\mu_I$ | Death rate for infected RBCs | 1/2 day<sup>-1</sup> | (59, 60) |
| $\mu_Z$ | Death rate for merozoites | 1/5 min<sup>-1</sup> | (65) |
| $m$ | Number of merozoites released per schizont | 15 | (60) |
| Structural Parameters |  |  |  |
| $n$ | Number of successive compartments for susceptible and for resistant RBCs | 25 | — |
| $n_I$ | Number of successive compartments for infected RBCs | 25 | — |

## Acknowledgments

This work was funded by NIH award 5R01HL139337 to MTD, and a Burroughs Wellcome Investigator in the Pathogenesis of Infectious Disease award #1016747 to COB. FC and EH were supported by award Number U54GM088558 from the National Institute Of General Medical Sciences. The content is solely the responsibility of the authors and does not necessarily represent the official views of the National Institute Of General Medical Sciences or the National Institutes of Health.

---

**Fig. S1.**
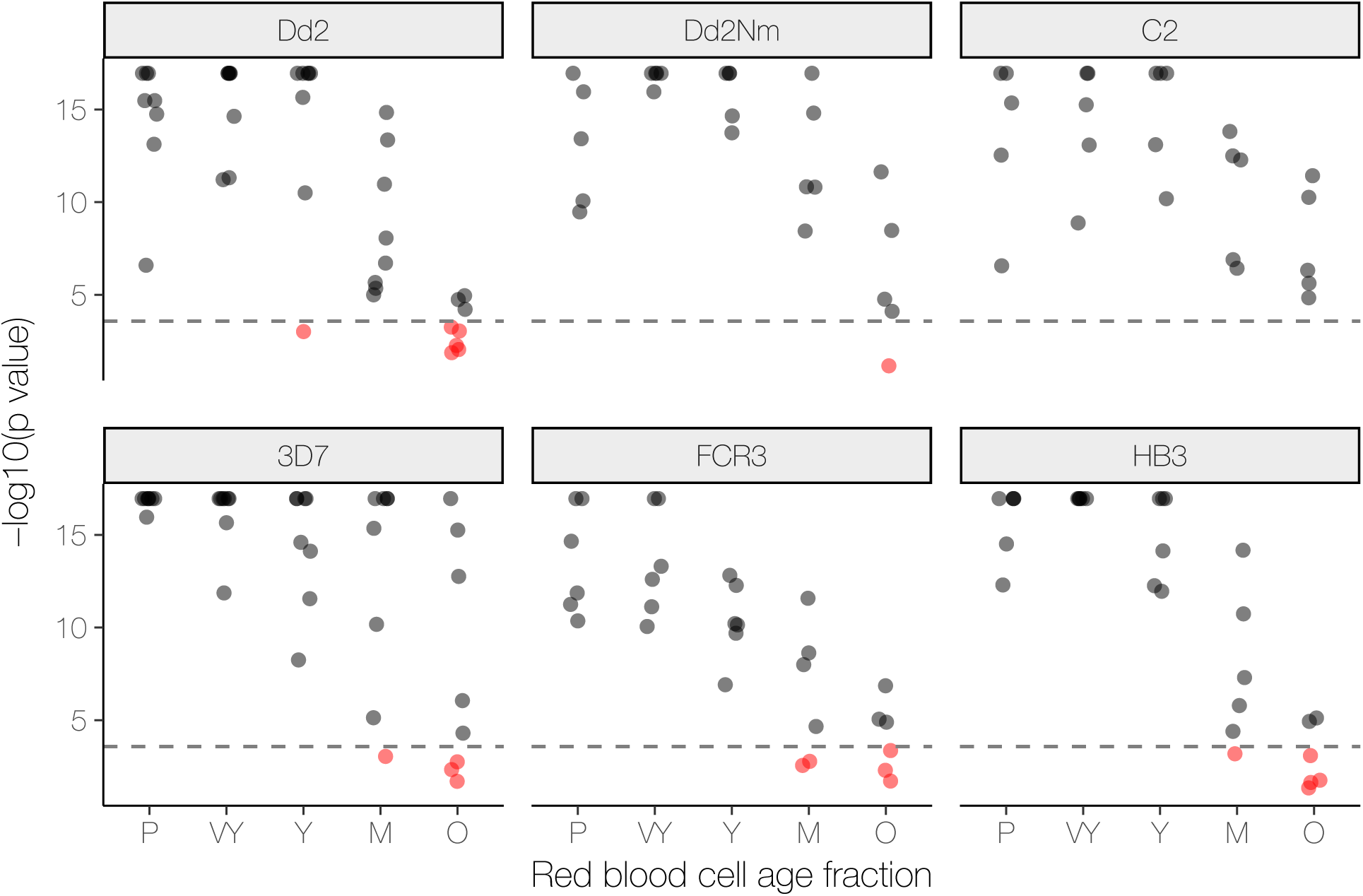
Comparison of Poisson and zero-inflated Poisson model fits. For each invasion assay, the empirical distribution of the number of parasites in a red blood cell was used to fit a Poisson model and a zero-inflated Poisson model. Model fits were compared using a likelihood ratio test. The p-values (N = 190, y-axis, negative log-transformed) are plotted above, organized by the parasite strain (panel title) and red blood cell age fraction (x-axis) used in the invasion assay. The abbreviations used in the x-axis are: P (pooled), VY (very young), Y (young), M (medium), O (old). For each combination of strain and age, multiple p-values, from different trials, are shown with their horizontal position jittered. The horizontal dashed lines in each panel are the significance cutoff assuming an overall significance level of 0.05 and Bonferroni correction. P-values lying below the line (highlighted in red) correspond to trials for which the zero-inflated Poisson model did not provide not a significantly better fit than the Poisson model.

**Fig. S2.**
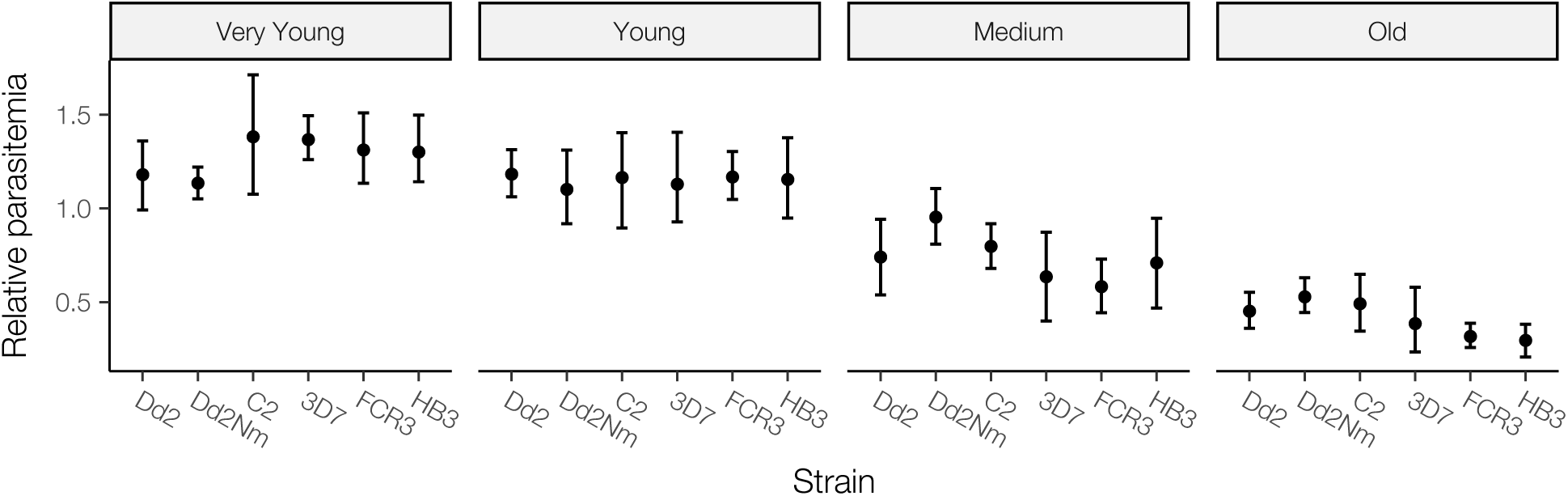
Relative post-invasion parasitemia versus strains. The distribution (mean with 95% bootstrap CI) of post-invasion parasitemia in each age fraction relative to pooled blood. The distributions are organized by strain (x-axis) and age fraction (panel title). For all age fractions, the Kruskal-Wallis rank sum test did not detect significant heterogeneity between strains (from youngest to oldest, p = 0.42, 0.94, 0.47, and 0.09).

**Fig. S3.**
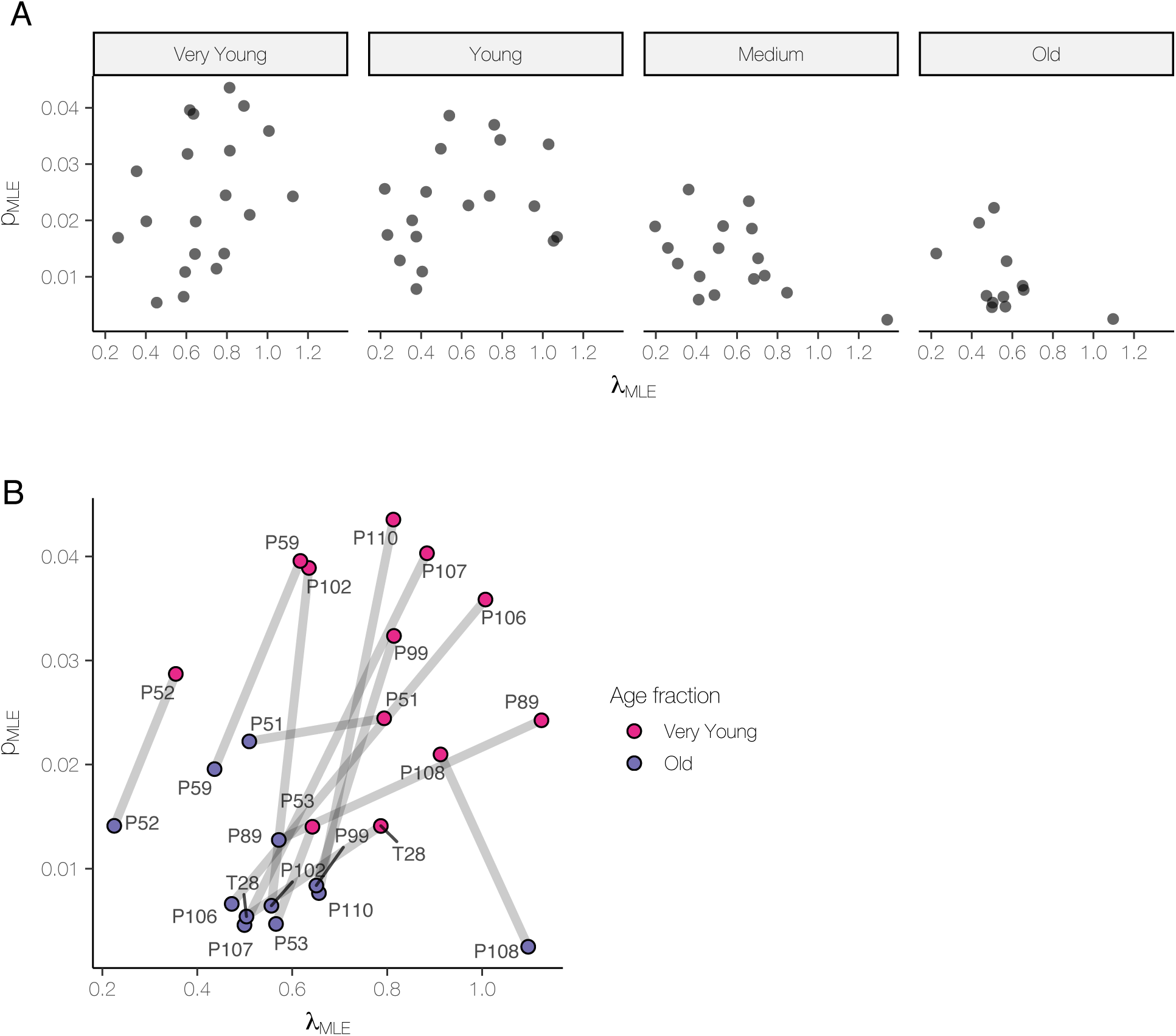
Invasion strategies of field strains. **(A)** Points show the maximum likelihood estimate of the fraction susceptible ***f***_***MLE***_ (y-axis) and ***λ***_***MLE***_ (x-axis) for field strains cultured *ex vivo* in each of four red blood cell age fractions (panel titles). For some trials, the model could not be fit due to small numbers of infected cells; there are 20, 18, 16, and 12 strains shown in the very young, young, medium, and old panels. **(B) *f***_***MLE***_ (y-axis) and ***λ***_***MLE***_ (x-axis) from (A) for the very young (pink) and old (blue) red blood cell age fractions are paired by strain here to highlight strain-specific responses to red blood cell aging.

**Fig. S4.**
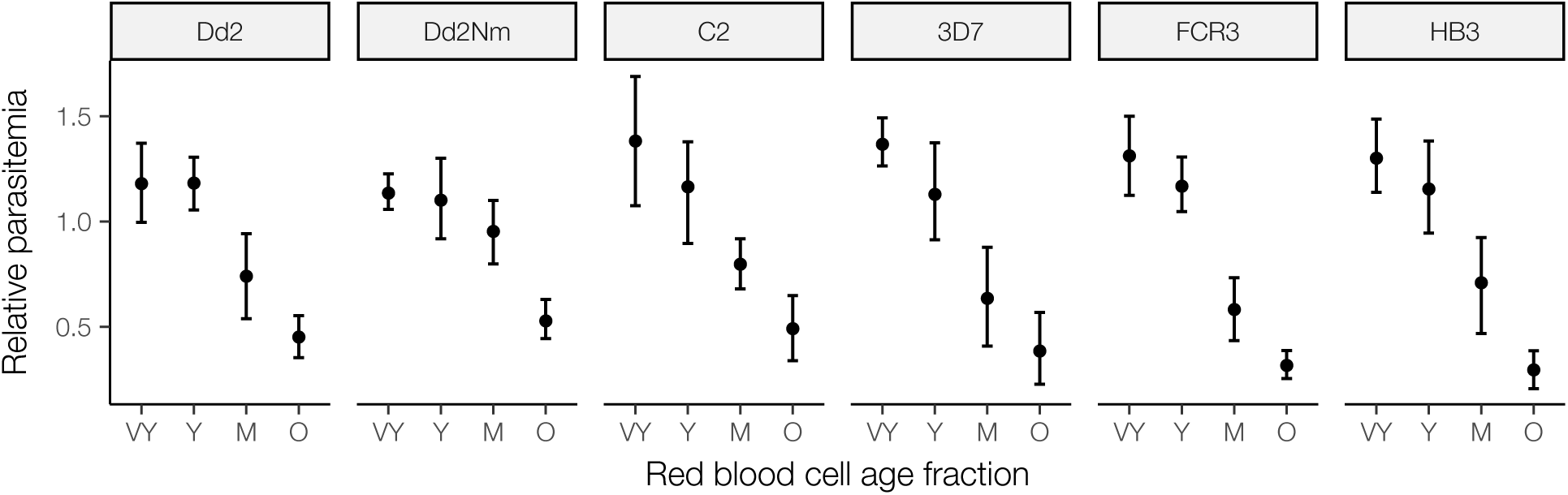
Relative post-invasion parasitemia versus age fractions. The distribution (mean with 95% bootstrap CI) of post-invasion parasitemia in each age fraction (x-axis) relative to pooled blood, organized by strain (panel title). The abbreviations used in the x-axis are: VY (very young), Y (young), M (medium), O (old). For each strain, the Kruskal-Wallis rank sum test detected significant heterogeneity between age fractions (from left to right: p < 0.01 for all strains), using an overall significance level of 0.05 with Bonferroni correction.

## References

1. Phillips A, Bassett P, Szeki S (2009) Risk factors for severe disease in adults with falciparum malaria. … Infectious Diseases.

2. Tangpukdee N, Krudsood S, Kano S, Wilairatana P (2012) Falciparum malaria parasitemia index for predicting severe malaria. Int J Lab Hematol 34(3):320–327.

3. Cowman AF, Crabb BS (2006) Invasion of red blood cells by malaria parasites. Cell 124(4):755–766.

4. Cortés A, et al. (2007) Epigenetic silencing of Plasmodium falciparum genes linked to erythrocyte invasion. PLoS Pathog 3(8):e107.

5. Jiang L, et al. (2010) Epigenetic control of the variable expression of a Plasmodium falciparum receptor protein for erythrocyte invasion. Proc Natl Acad Sci USA 107(5):2224–2229.

6. Mayer DCG, et al. (2004) Polymorphism in the Plasmodium falciparum erythrocyte-binding ligand JESEBL/EBA-181 alters its receptor specificity. Proc Natl Acad Sci USA 101(8):2518–2523.

7. Triglia T, Duraisingh MT, Good RT, Cowman AF (2005) Reticulocyte-binding protein homologue 1 is required for sialic acid-dependent invasion into human erythrocytes by Plasmodium falciparum. Mol Microbiol 55(1):162–174.

8. Gaur D, Mayer DCG, Miller LH (2004) Parasite ligand-host receptor interactions during invasion of erythrocytes by Plasmodium merozoites. Int J Parasitol 34(13-14):1413–1429.

9. Allison AC (1954) Protection afforded by sickle-cell trait against subtertian malareal infection. British Medical Journal 1(4857):290–294.

10. Ruwende C, et al. (1995) Natural selection of hemi-and heterozygotes for G6PD deficiency in Africa by resistance to severe malaria. Nature 376(6537):246–249.

11. Genton B, et al. (1995) Ovalocytosis and cerebral malaria. Nature 378(6557):564–565.

12. Allen SJ, et al. (1997) alpha+-Thalassemia protects children against disease caused by other infections as well as malaria. Proc Natl Acad Sci USA 94(26):14736–14741.

13. Hutagalung R, Wilairatana P (1999) Influence of hemoglobin E trait on the severity of Falciparum malaria. Journal of infectious ….

14. Agarwal A, et al. (2000) Hemoglobin C associated with protection from severe malaria in the Dogon of Mali, a West African population with a low prevalence of hemoglobin S. Blood 96(7):2358–2363.

15. Maier AG, et al. (2003) Plasmodium falciparum erythrocyte invasion through glycophorin C and selection for Gerbich negativity in human populations. Nat Med 9(1):87–92.

16. Cockburn IA, et al. (2004) A human complement receptor 1 polymorphism that reduces Plasmodium falciparum rosetting confers protection against severe malaria. Proc Natl Acad Sci USA 101(1):272–277.

17. Gaur D, Chitnis CE (2011) Molecular interactions and signaling mechanisms during erythrocyte invasion by malaria parasites. Curr Opin Microbiol 14(4):422–428.

18. Gunalan K, Gao X, Yap SSL, Huang X, Preiser PR (2013) The role of the reticulocyte-binding-like protein homologues of Plasmodium in erythrocyte sensing and invasion. Cell Microbiol 15(1):35–44.

19. Tham W-H, Healer J, Cowman AF (2012) Erythrocyte and reticulocyte binding-like proteins of Plasmodium falciparum. Trends in Parasitology 28(1):23–30.

20. Crosnier C, et al. (2011) Basigin is a receptor essential for erythrocyte invasion by Plasmodium falciparum. Nature 480(7378):534–537.

21. Shinozuka T (1994) Changes in human red blood cells during aging in vivo. Keio J Med 43(3):155–163.

22. Aminoff D (1988) The role of sialoglycoconjugates in the aging and sequestration of red cells from circulation. Blood Cells 14(1):229–257.

23. Lutz HU, Fehr J (1979) Total sialic acid content of glycophorins during senescence of human red blood cells. J Biol Chem 254(22):11177–11180.

24. Simpson JA, Silamut K, Chotivanich K, Pukrittayakamee S, White NJ (1999) Red cell selectivity in malaria: a study of multiple-infected erythrocytes. Trans R Soc Trop Med Hyg 93(2):165–168.

25. McKenzie FE, Jeffery GM, Collins WE (2002) PLASMODIUM VIVAX BLOOD-STAGE DYNAMICS. J Parasitol 88(3):521–535.

26. Pasvol G, Weatherall DJ, Wilson RJ (1980) The increased susceptibility of young red cells to invasion by the malarial parasite Plasmodium falciparum. Br J Haematol 45(2):285–295.

27. Iyer J, Grüner AC, Rénia L, Snounou G, Preiser PR (2007) Invasion of host cells by malaria parasites: a tale of two protein families. Mol Microbiol 65(2):231–249.

28. Tham W-H, et al. (2010) Complement receptor 1 is the host erythrocyte receptor for Plasmodium falciparum PfRh4 invasion ligand. Proc Natl Acad Sci USA 107(40):17327–17332.

29. Dolan SA, Miller LH, Wellems TE (1990) Evidence for a switching mechanism in the invasion of erythrocytes by Plasmodium falciparum. J Clin Invest 86(2):618–624.

30. Reed MB, et al. (2000) Targeted disruption of an erythrocyte binding antigen in Plasmodium falciparum is associated with a switch toward a sialic acid-independent pathway of invasion. Proc Natl Acad Sci USA 97(13):7509–34.

31. Duraisingh MT, Maier AG, Triglia T, Cowman AF (2003) Erythrocyte-binding antigen 175 mediates invasion in Plasmodium falciparum utilizing sialic acid-dependent and -independent pathways. Proc Natl Acad Sci USA 100(8):4796–4801.

32. Deans A-M, et al. (2007) Invasion pathways and malaria severity in Kenyan Plasmodium falciparum clinical isolates. Infect Immun 75(6):3014–3020.

33. Bowyer PW, et al. (2015) Variation in Plasmodium falciparum erythrocyte invasion phenotypes and merozoite ligand gene expression across different populations in areas of malaria endemicity. Infect Immun 83(6):2575–2582.

34. Baum J, Pinder M, Conway DJ (2003) Erythrocyte invasion phenotypes of Plasmodium falciparum in The Gambia. Infect Immun 71(4):1856–1863.

35. Baum J, et al. (2009) Reticulocyte-binding protein homologue 5 - an essential adhesin involved in invasion of human erythrocytes by Plasmodium falciparum. Int J Parasitol 39(3):371–380.

36. Simpson JA, Aarons L, Collins WE, Jeffery GM, White NJ (2002) Population dynamics of untreated Plasmodium falciparum malaria within the adult human host during the expansion phase of the infection. Parasitology 124(Pt 3):247–263.

37. Chotivanich K, et al. (2000) Parasite multiplication potential and the severity of Falciparum malaria. J Infect Dis 181(3):1206–1209.

38. Deans A-M, et al. (2006) Low multiplication rates of African Plasmodium falciparum isolates and lack of association of multiplication rate and red blood cell selectivity with malaria virulence. Am J Trop Med Hyg 74(4):554–563.

39. Stubbs J, et al. (2005) Molecular mechanism for switching of P. falciparum invasion pathways into human erythrocytes. Science 309(5739):1384–7.

40. Duraisingh MT, et al. (2004) Phenotypic variation of Plasmodium falciparum merozoite proteins directs receptor targeting for invasion of human erythrocytes. EMBO J 22(5):1047–57.

41. Persson KEM, et al. (2013) Erythrocyte-binding antigens of Plasmodium falciparum are targets of human inhibitory antibodies and function to evade naturally acquired immunity. J Immunol 191(2):785–794.

42. Persson KEM, et al. (2008) Variation in use of erythrocyte invasion pathways by Plasmodium falciparum mediates evasion of human inhibitory antibodies. J Clin Invest 118(1):342–351.

43. Lim C, Hansen E, DeSimone TM, Moreno Y (2013) Expansion of host cellular niche can drive adaptation of a zoonotic malaria parasite to humans. Nature.

44. Kerlin DH, Gatton ML (2013) Preferential invasion by Plasmodium merozoites and the selfregulation of parasite burden. PLoS ONE 8(2):e57434.

45. McQueen PG, McKenzie FE (2004) Age-structured red blood cell susceptibility and the dynamics of malaria infections. Proc Natl Acad Sci USA 101(24):9161–9166.

46. Hyman JM, Li J (2000) An intuitive formulation for the reproductive number for the spread of diseases in heterogeneous populations. Math Biosci 167(1):65–86.

47. Cohen NS, Ekholm JE, Luthra MG, Hanahan DJ (1976) Biochemical characterization of density-separated human erythrocytes. Biochim Biophys Acta 419(2):229–242.

48. Rennie CM, Thompson S, Parker AC, Maddy A (1979) Human erythrocyte fraction in “Percoll” density gradients. Clin Chim Acta 98(1-2):119–125.

49. Romero PJ, Romero EA, Winkler MD (1997) Ionic calcium content of light dense human red cells separated by Percoll density gradients. Biochim Biophys Acta 1323(1):23–28.

50. Prall YG, Gambhir KK, Ampy FR (1998) Acetylcholinesterase: an enzymatic marker of human red blood cell aging. Life Sci 63(3):177–184.

51. Omodeo-Salè F, et al. (2003) Accelerated senescence of human erythrocytes cultured with Plasmodium falciparum. Blood 102(2):705–711.

52. Bratosin D, et al. (1998) Cellular and molecular mechanisms of senescent erythrocyte phagocytosis by macrophages. A review. Biochimie 80(2):173–195.

53. (null) RCT (2017) R: A Language and Environment for Statistical Computing (R Foundation for Statistical Computing, Vienna, Austria) Available at: https://www.R-project.org/.

54. Dinno A dunn.test: Dunn’s Test of Multiple Comparisons Using Rank Sums. Available at: https://CRAN.R-project.org/package=dunn.test.

55. Bianconi E, et al. (2013) An estimation of the number of cells in the human body. Ann Hum Biol 40(6):463–471.

56. Berlin NI, Waldmann TA, Weissman SM (1959) Life span of red blood cell. Physiological Reviews 39(3):577–616.

57. Frevert U (2004) Sneaking in through the back entrance: the biology of malaria liver stages. Trends in Parasitology 20(9):417–424.

58. Rosenberg R, Wirtz RA, Schneider I, Burge R (1990) An estimation of the number of malaria sporozoites ejected by a feeding mosquito. Trans R Soc Trop Med Hyg 84(2):209–212.

59. Beier JC, Davis JR, Vaughan JA, Noden BH, Beier MS (1991) Quantitation of Plasmodium falciparum sporozoites transmitted in vitro by experimentally infected Anopheles gambiae and Anopheles stephensi. Am J Trop Med Hyg 44(5):564–570.

60. Beier MS, Davis JR, Pumpuni CB, Noden BH, Beier JC (1992) Ingestion of Plasmodium falciparum sporozoites during transmission by anopheline mosquitoes. Am J Trop Med Hyg 47(2):195–200.

61. Trager W, Jensen JB (1976) Human malaria parasites in continuous culture. Science 193(4254):673–675.

62. Reilly HB, Wang H, Steuter JA, Marx AM, Ferdig MT (2007) Quantitative dissection of clone-specific growth rates in cultured malaria parasites. Int J Parasitol 37(14):1599–1607.

63. Lloyd AL (2001) Destabilization of epidemic models with the inclusion of realistic distributions of infectious periods. Proc Biol Sci 268(1470):985–993.

64. Baer K, Klotz C, Kappe SHI, Schnieder T, Frevert U (2007) Release of hepatic Plasmodium yoelii merozoites into the pulmonary microvasculature. PLoS Pathog 3(11):e171.

65. Miller LH, Ackerman HC, Su X-Z, Wellems TE (2013) Malaria biology and disease pathogenesis: insights for new treatments. Nat Med 19(2):156–167.

66. Crick AJ, et al. (2013) An automated live imaging platform for studying merozoite egress-invasion in malaria cultures. Biophys J 104(5):997–1005.

67. Boyle MJ, et al. (2010) Isolation of viable Plasmodium falciparum merozoites to define erythrocyte invasion events and advance vaccine and drug development. Proc Natl Acad Sci USA 107(32):14378–14383.

